# Correlation between Retinal Ganglion Cell Loss and Nerve Crush Force-Impulse Established with Instrumented Tweezers in Mice

**DOI:** 10.1101/687616

**Authors:** Xiaorong Liu, Liang Feng, Ishan Shinde, James D. Cole, John B. Troy, Laxman Saggere

## Abstract

**Objectives:** Rodent models of optic nerve crush (ONC) have often been used to study degeneration and regeneration of retinal ganglion cells (RGCs) and their axons as well as the underlying molecular mechanisms. However, ONC results from different laboratories exhibit a range of RGC injury with varying degree of axonal damage. We developed an instrumented tweezers to measure optic nerve (ON) crush forces in real time and studied the correlation between RGC axon loss and force-impulse, the product of force and duration, applied through the instrumented tweezers in mice.

**Methods:** A pair of standard self-closing #N7 tweezers were instrumented with miniature foil strain gauges at optimal locations on both tweezer arms. The instrumented tweezers were capable of recording the tip closure forces in the form of voltages, which were calibrated through load cells to corresponding tip closure forces over the operating range. Using the instrumented tweezers, the ONs of multiple mice were crushed with varied forces and durations and the axons in the immunostained sections of the crushed ONs were counted.

**Results:** We found that the surviving axon density correlated with crush force, with longer duration and stronger crush forces producing consistently more axon damage.

**Discussion:** The instrumented tweezers enable a simple technique for measurement of ONC forces in real-time for the first time. Using the instrumented tweezers, experimenters can quantify crush forces during ONC to produce consistent and predictable post-crush cell death. This should permit future studies a way to produce nerve damage more consistently than is available now.

## Introduction

Rodent models of optic nerve crush (ONC) have often been used to study degeneration and regeneration of retinal ganglion cells (RGCs) and the underlying molecular mechanisms [1–8]. For example, RGC death may be triggered by different signaling paths in different forms of glaucoma, so experimental models have been developed to mimic these different forms. Although an experimental ONC injury may not be a “true” model of glaucoma, it represents an optic neuropathy leading to acute RGC death that is often assumed to involve some of the same molecular mechanisms as those responsible for RGC death in certain types of glaucoma. Moreover, rodent models of ONC offer the advantage of producing a rapid and profound loss of RGCs that expedite investigations of neural regeneration [9–11].

However, a major drawback with current rodent models of ONC is that the degree of RGC loss for each crush trial seems to vary substantially from laboratory to laboratory (Table 1) and, within a laboratory, is observed to vary from investigator to investigator (unpublished data from Liu laboratory). Some of the seeming variance in RGC loss can be attributed to methodological differences (see Notes in Table 1), but some of the variance cannot. For example, at 2-weeks (w) following ONC, Leung et al. [12] reported about 89% RGC loss, Duan et al. [4] found about 80% RGC loss, while our results indicated about 70% RGC loss [13]. At a slightly shorter time period (9-days) after crush, Kalesnykas et al. [2] reported about 98% of RGC axons died and RGC number decreased by 64%. All of these studies employed methods in which RGC or optic nerve (ON) axon loss was quantified reliably. The extent to which these differences can be ascribed to variation between laboratories and variation between individuals performing the crush is unclear, although RGC or axon loss is likely relatively consistent within each laboratory when one person performs the crush; e.g., individuals from our laboratory achieve less than ±12% variation (standard deviation) in measurements of RGC loss from experiment to experiment [7,13,14].

**Table 1.**
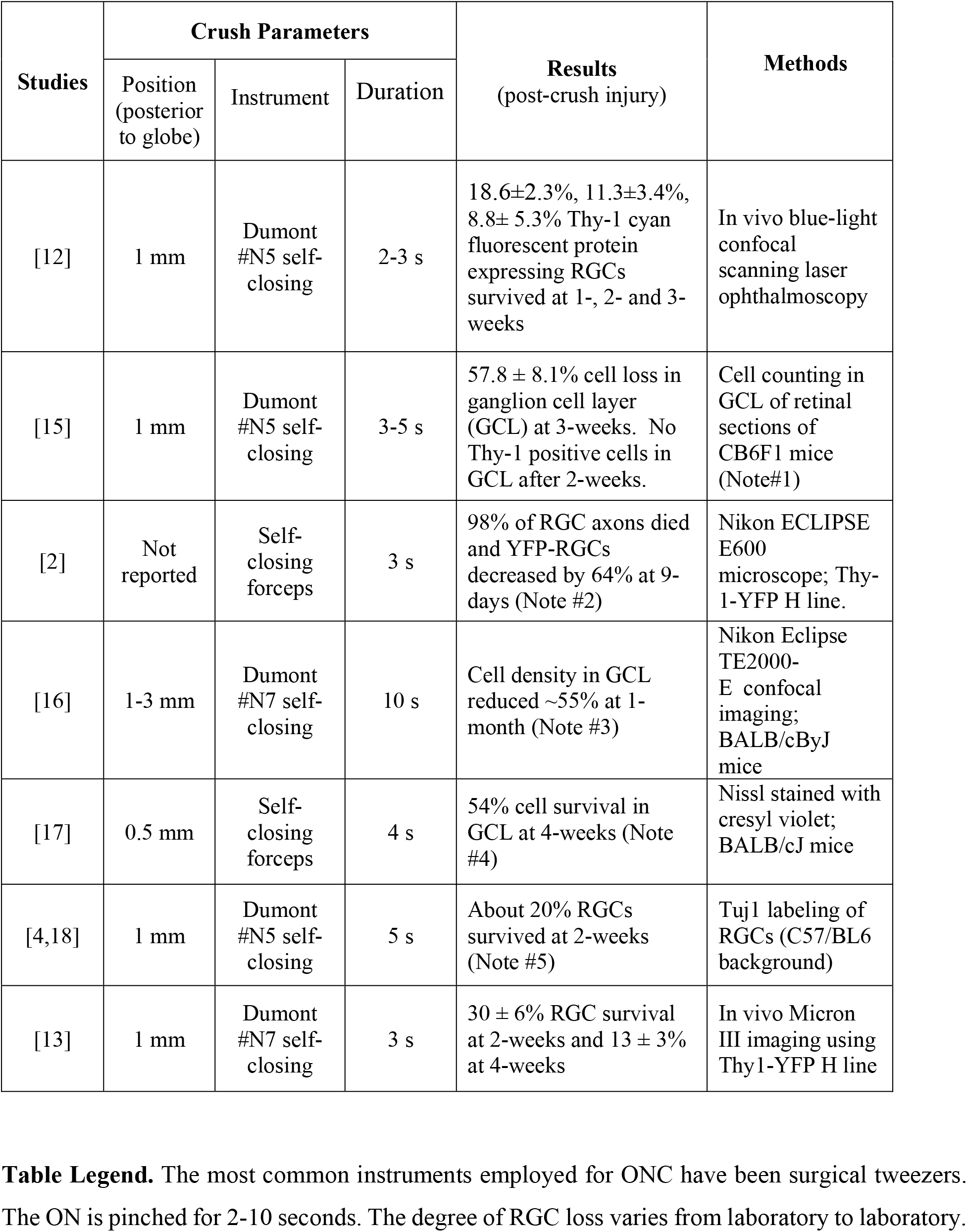
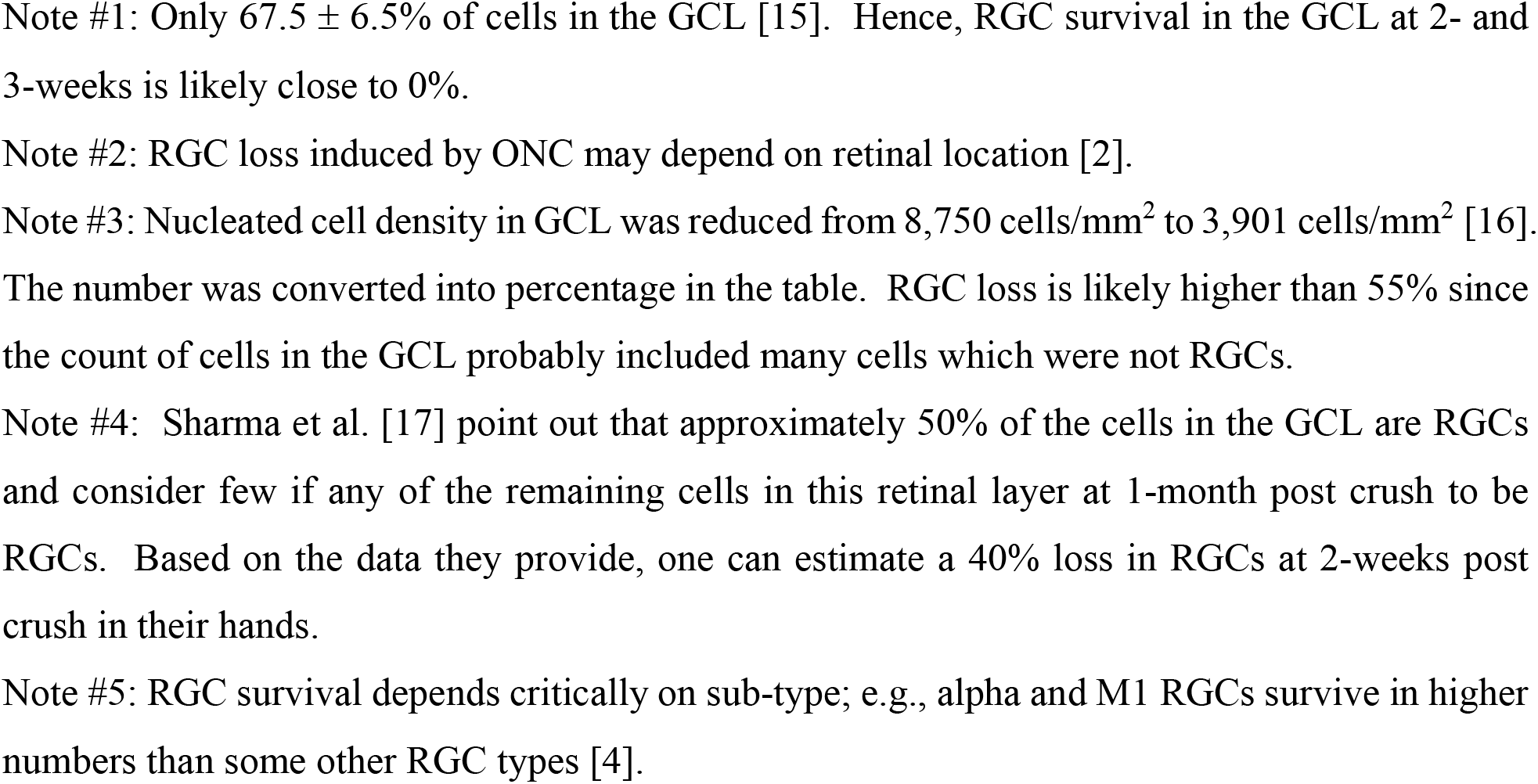
Comparison of crush setups and corresponding RGC loss from different studies.

The inter-laboratory variability in ONC models could be reduced if a standard calibrated tool for crushing were available. The common crush technique (pinching the nerve with a tool for 2-3 seconds) and the resulting damage rely on manual actuation of the tool and operator characteristics of holding the tool during an ONC trial. Obviously, this approach is subject to variation from investigator to investigator. As a result, the pattern of the axonal damage can vary, making comparison between different groups difficult, thus reducing the value of this technique for studies of the pathways underlying injury-induced axonal degeneration and RGC death. Understanding these pathways well is clearly a prerequisite for developing therapies to circumvent cell loss and encourage axonal regeneration following ON injury.

In this study, we examined how varying duration and force together affects RGC survival following an ONC injury in mice using instrumented tweezers with the objective of identifying a means to better control the degree of nerve injury.

## Material and Methods

### Development of Instrumented Tweezers

First, the mechanical strains experienced by the tweezers during its normal range of operation was analyzed via Finite Element Analysis in COMSOL Multiphysics software, where the exact geometry of the tweezers including their curved tips (0.03×0.07 mm^2^), obtained through imaging, and the tweezers’ material properties, assumed to be INOX stainless steel with the Young’s modulus of 200 GPa, were represented. Based on the simulation and analysis of the mechanics of the tweezers, the mechanical strain was found to range from 0 to 10^-4^ corresponding to a maximum tip displacement of 5.4 mm, with the maximum strain located at a point approximately 16 mm from the base of the tweezers.

We evaluated several different types of sensors including foil strain gauges, piezoelectric sensors, polyvinylidene fluoride thin films, optical fiber Bragg gratings and custom MEMS-based sensors and weighed against desirable sensor characteristics such as sensitivity, resolution, bandwidth, signal-to-noise ratio, voltage requirements, robustness, portability packaging and the area available for mounting sensors on the tweezers. The commercially available miniature precision foil strain gauge (SGD-3/120-LY11, Omega Engineering, USA) with an overall footprint of 3 mm × 3.8 mm was found to be best suited for integration with the tweezers in this study.

Two miniature precision foil strain gauges were mounted on the #N7 tweezers, one on each arm centered 16 mm from the base of the tweezers for maximum signal sensitivity and packaged using Parafilm M® to provide electrical insulation, water resistance and mechanical protection. The leads from each strain gauge were soldered to 32 American wire gauge wires and encapsulated in a heat shrink tube for safety and portability. The other ends of the lead wires were connected to a data acquisition (DAQ) system using a three wire quarter bridge configuration via push-in connectors (Figure 1). The DAQ system comprised of a bridge completion module, a voltage source and an analog-to-digital DAQ board (USB-6212 BNC, National Instruments Corp., USA), with the strain gauge data acquired by LabVIEW software [19].

**Figure 1.**
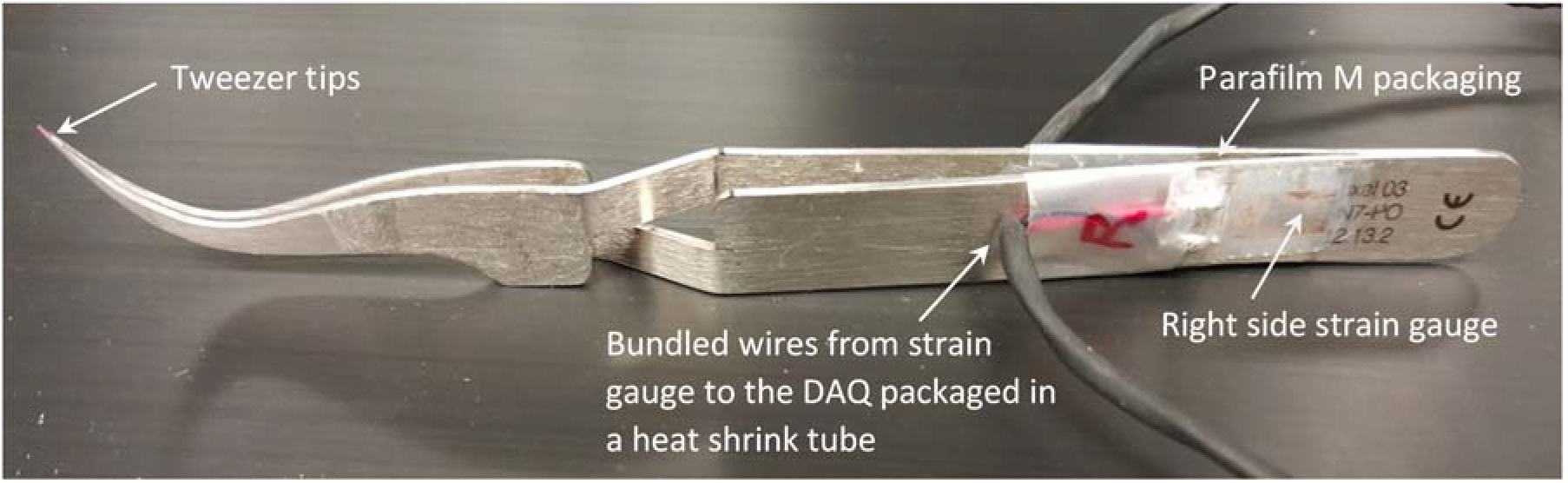
Cross-action Dumont #N7 tweezers instrumented with miniature precision foil strain gauges and packaged to protect against damage due to mechanical handling.

The compressive tip forces for various openings of tweezer tips and the corresponding strains were measured by placing a subminiature load cell with 1000 g load range (Model 13, Honeywell Inc., USA) between the tips of the tweezers and recording both load cell signals and voltage signals from the strain gauges while the tweezers were held firmly fixed in a clamp at the base for tip openings ranging from 3.3 mm (the thickness of the load cell) to 5.5 mm in increments of 0.25 mm. The input actuation force that an individual would apply to push open the tweezers was also measured via another load cell held in contact with the tweezers near the bridge area and advancing a micrometer spindle from 0 to 1.2 mm in increments of 50 μm to push on the tweezer arms. Figure 2 shows a schematic of the experiment setup for force and strain measurements during calibration. The incremental opening of the tweezer tips from 3.3 mm to 5.5 mm was accomplished by inserting standard thickness feeler gauges between the tips and the load cell. For tweezer tip openings from 0 to 3.3 mm, accomplished by inserting feeler gauges between the tweezer tips, only the strains were measured. The data from the two sets of measurements (for tip openings from 0 to 3.3 mm and 3.3 to 5.5 mm), combined through a regression analysis, were used to establish the relationship between compressive forces of the tweezer tips and the corresponding strains for tip openings from 0 to 5.5 mm [19].

**Figure 2.**
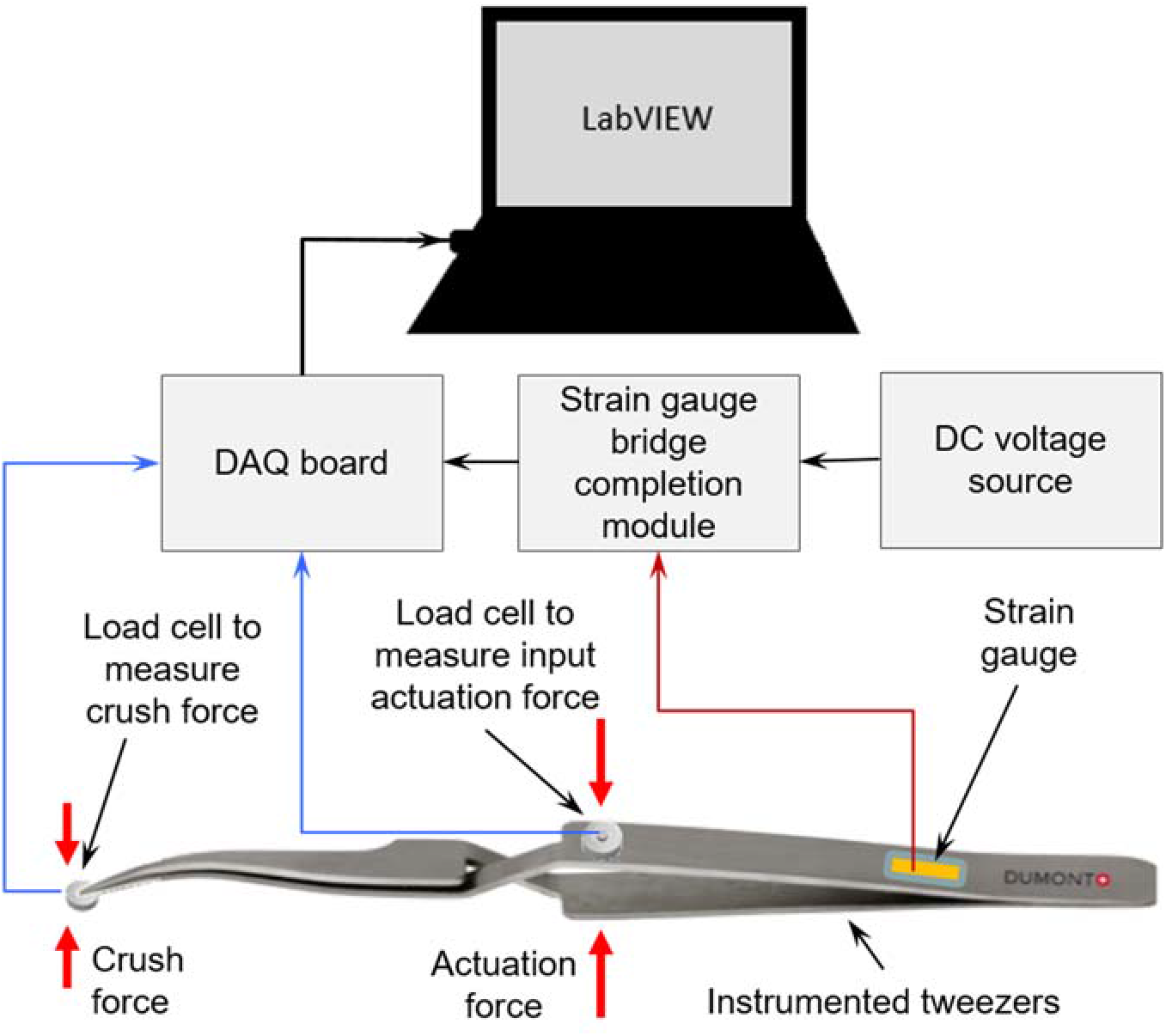
Schematic of the experimental setup used for measurement of forces and strains during calibration of the instrumented tweezers. The DAQ system comprised a DAQ board, a strain gauge bridge completion module and a DC voltage supply. A micrometer spindle (not shown in the schematic) was used to deliver the input actuation force and push on the tweezer arms.

### Optic Nerve Crush Injury and Axon Counting

Mice were anesthetized through intraperitoneal injection of ketamine (100 mg/kg) as described previously [13,7]. In brief, small cuts were made in the superior and lateral conjunctiva to expose the ON. The ONs were crushed with the instrumented #N7 reverse grip tweezers at a distance between 0.5 and 1 mm from the globe. Both the intensity of the crush force applied and the duration of applied force in each trial were recorded through the data acquisition system connected to the instrumented tweezers. In each crush trial, the strain gauges were first normalized and their initial readings were reset to zero, and then, strain output signals from the instrumented tweezers were continuously acquired by the DAQ system from the moment the operator grasped the ON between the tweezers’ tips until the ON was released from the tweezers’ grasp. The duration of crush was varied from 2 to 8 seconds, which covers the common time range used by various laboratories. Finally, the strains recorded in the form of voltages were scaled to corresponding crush forces, according to the calibration.

Mice were sacrificed at one or three days following crush, and their ONs dissected, cryo-sectioned, and immunostained for axon counting. Cryo-Sections (14-18 μm thick) were taken between 0.5 and 1 mm from the ON head and stained using p-Phenylenediamine (PPD 1:50, Fisher). The stained sections were then imaged using a Zeiss Observer A1 microscope (Carl-Zeiss, Oberkochen, Germany) and a total of 20 images covering the center, mid-periphery, and peripheral margin of each ON were recorded. The axons in each of the three regions of the ON were counted using Imaris software (Bitplane, Concord, MA). Counts were averaged and normalized with respect to the area of the region of ON studied to obtain axon density [20].

All animal experiments were performed in the Liu laboratory at Northwestern University. All animal procedures were approved by the Northwestern University Institutional Animal Care and Use Committee and performed in accordance with the guidelines on the Use of Animals in Neuroscience Research from the National Institutes of Health and the Society for Neuroscience.

## Results

To measure the compressive forces between the tips of a self-closing #N7 tweezer during its normal range of operation, we selected miniature precision foil strain gauges (SGD-3/120-LY11, Omega Engineering, USA) and mounted one strain gauge on each arm of the tweezers (Figure 1). When an object is held between the tips of the tweezers, the strain gauges mounted on the tweezer arms transduce the compressive force applied by the tweezers on the object into voltage signals (Figure 2). Following the calibration as described in the Methods, the experimenter was trained to maneuver and actuate the tweezers expertly. Next, crush forces were applied for various amounts of time to induce injury whose magnitude was measured through RGC survival.

To investigate whether crush duration might affect the force intensity delivered by an experimenter or *vice versa,* we examined if these parameters might be correlated. They were not. Figure 3 plots intensity *versus* duration for all ONC trials performed by a single experimenter undertaken for this study. Strong or weak crush forces were delivered each time randomly and they showed no correlation with duration.

**Figure 3.**
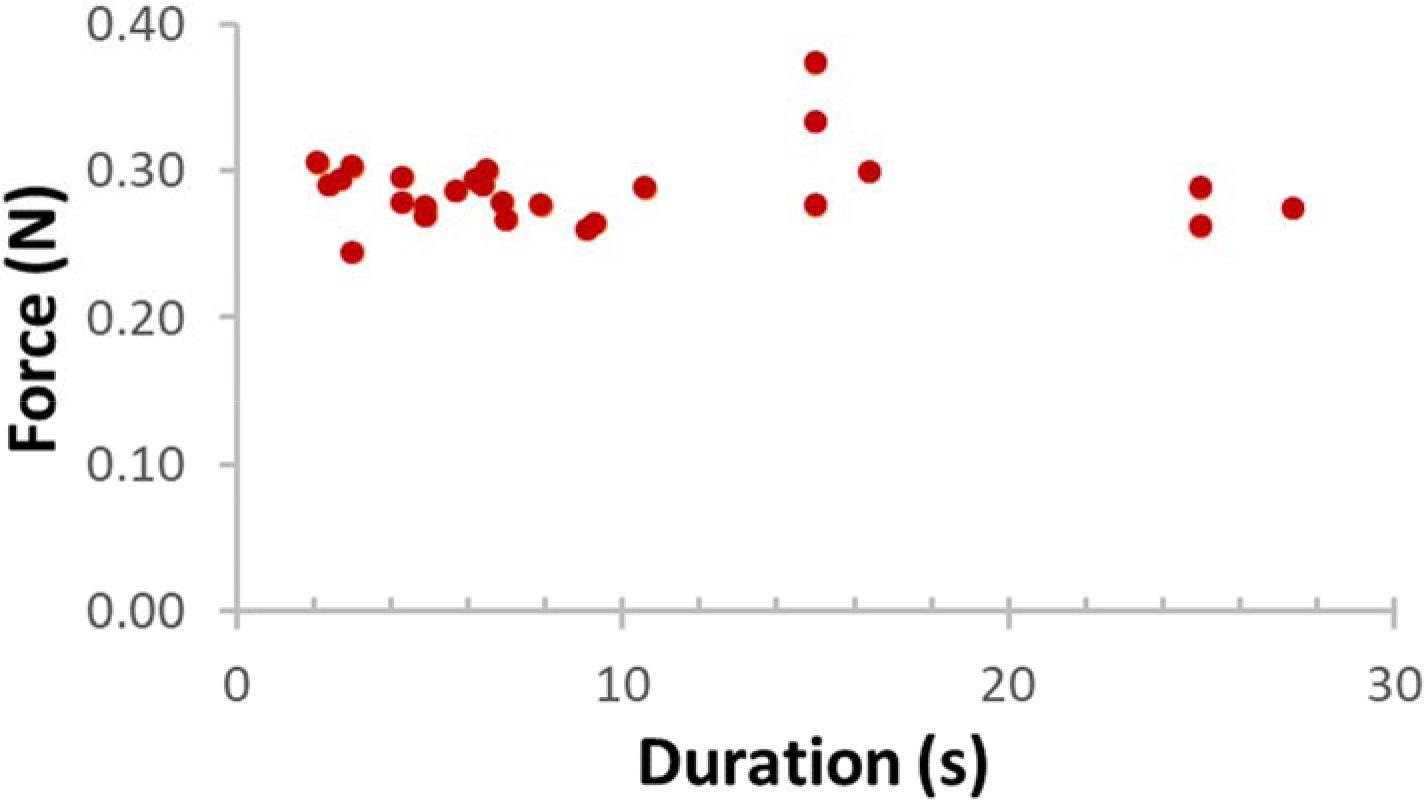
Plot of crush forces delivered using the instrumented self-closing tweezer by a single experimenter for various durations during ONC trials. The data show no correlation between the duration of the crush and force delivered. The data points scatter around a mean force of 0.287 N with a standard deviation of 0.025 N. With the instrumented self-closure tweezers, we sought to deliver a consistent force with each trial. With the occasional outlier, it is evident from Figure 3 that a single investigator can operate the Dumont #N7 self-closing tweezers to generate a largely consistent crush force with just ±8.7% variation (standard deviation) irrespective of duration.

As shown in Figure 4, ONC produces known signs of axon degeneration and loss [21,9,20]. At one-day post-ONC injury, there was a significant loss of axon density (Figure 4). In control mice, individual axons (dark PPD stained circles) appear normal and tightly packed, while crushed ONs have fewer axons per section. Additionally, the sections reveal other signs of degeneration, most notably the accumulation of vacuoles amongst dead or dying axons (orange arrow, Figure 4D), axonal swelling (green arrows, Figure 4D), and the abundance of degenerative debris in the form of dystrophic neurites (blue arrows, Figure 4D).

**Figure 4.**
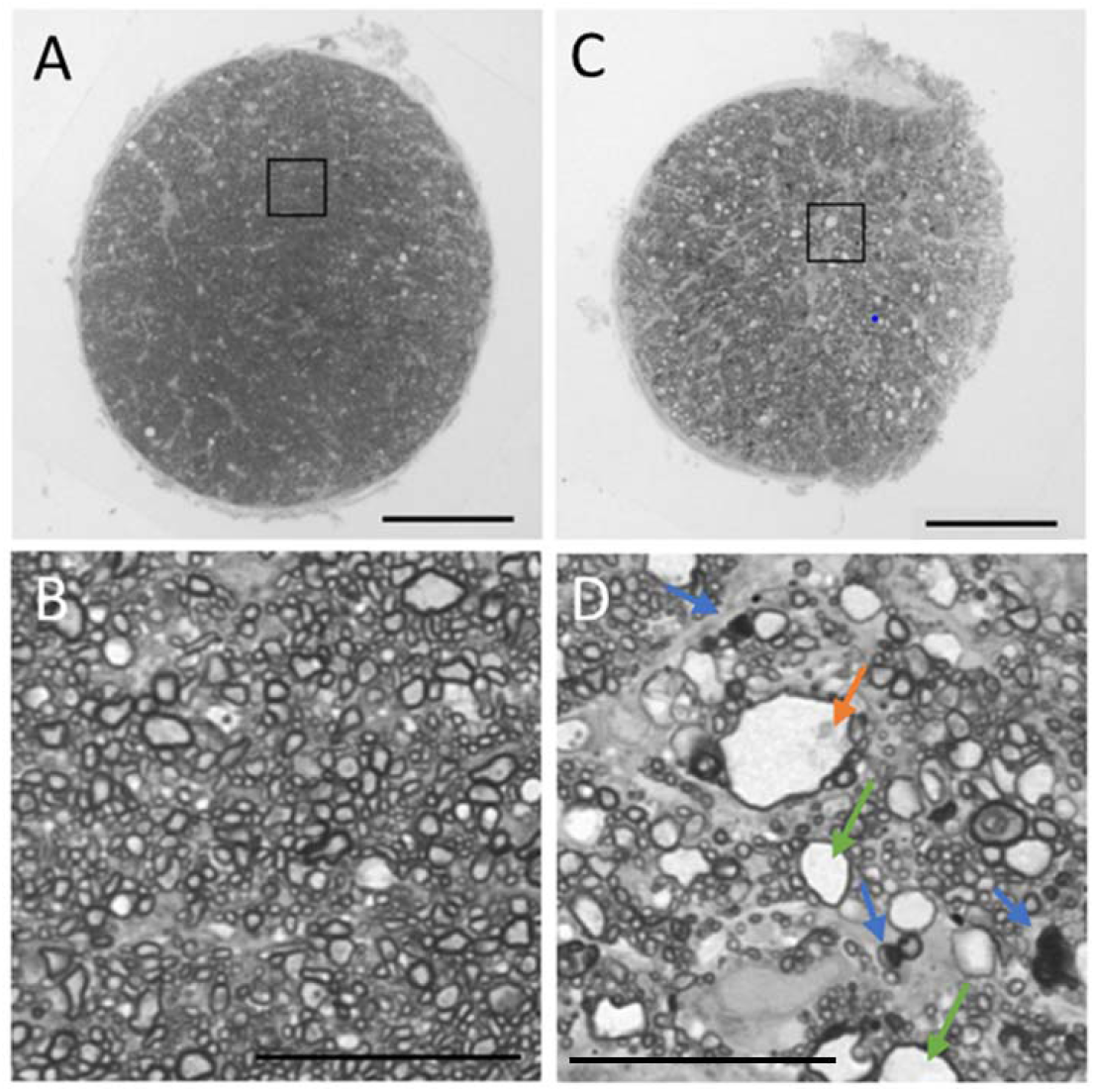
Microscope images of the control and crushed ON cross sections studied. (A) Cross-section of the uncrushed ON (control). (B) Higher magnification image of the black-bordered square region marked on A. (C) Cross-section of the one-day post-crush ON. (D) Higher magnification image of the black-bordered square region marked on C. In all images, each dark-bordered irregular circle represents one axon immunostained by PPD (1:50, Fisher). After ONC, the axons exhibited degenerative signs such as decreased axon density, widespread vacuole accumulation (orange arrow, image D), axon swelling (green arrows, image D), and prevalence of degenerative debris such as the dystrophic neurites (blue arrows, image D). (A, C) Scale bar = 100 μm and (B, D) Scale bar = 50 μm.

To study the combined effects of the crush duration and the crush force, we plotted axon density against the crush force impulse (newton-second), which revealed that axon density declines with increased force impulse (Figure 5). This relationship is represented with a consistent and predictable power trend line (R^2^ = 0.97), reaching an asymptotic density as force-impulse approaches 8 newton-second. Our results indicate that prolonged crush duration and increased force leads to more axon degeneration. Most interestingly, the line of best fit predicts the dependence of axon density upon force impulse over the range studied with a deviation of data points from the line of just 4.4% (standard deviation).

**Figure 5.**
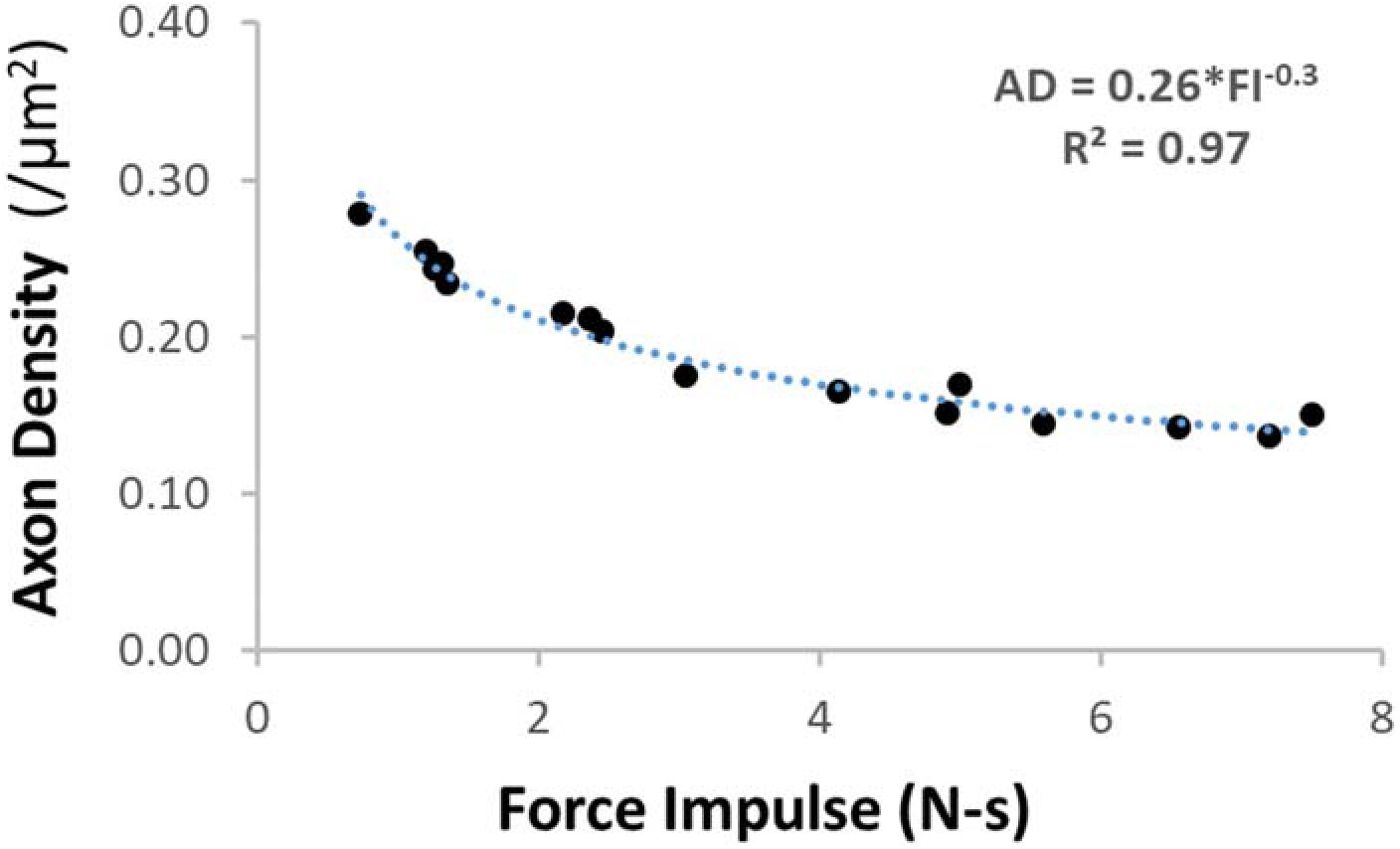
Plot of axon density of a 3-day post-crush ON plotted against the force impulse (newton-second). The resulting curve indicates that prolonged ONC and increased force of that crush will lead to decreased axon density at 3-days post-crush. AD: axon density; FI: force impulse.

## Discussion

Historically, a variety of tools and methods include silk threads [22], micro-slings [23], clamps [24], aneurysm or surgical clips [25–28], inflatable cuffs [29,30], custom microinjuring devices [31,32], and different types of forceps (shown in Table 1) have been used by various groups to study RGC death induced by the ONC injury in different animal models. Using different types of force measurement techniques, many of these groups have induced graded ONC and quantified the severity of the ONC, with an ultimate goal of creating a standardized crush model. For example, Burke et al. [29] and Cottee et al. [30] used a deflated balloon fastened inside a brass cylinder, which was hooked around the ON via an extensive surgical procedure in a cat model. The ON was squeezed by inflating the balloon against the inside of the cylinder. They reported that the pressure required to produce an effective nerve block ranged from 70 to 200 kPa but that pressure had no absolute significance since the balloon did not closely envelop the nerve at zero pressure [30]. Gellrich et al. [31] used a micro-hook mounted on the tip of a small dynamometer maneuvered by a conventional micromanipulator to deliver graded crush forces to the ON in a rat model. They reported that neuron death in the RGC layer directly correlated with both the crush pressures (5-20 cN/cm^2^) and durations (1-30 seconds). Klöcker et al. [32] also used a similar setup for ONC in a rat model and reported that RGC loss was significant with forces ≥ 10 cN under the assumption that the ON flattened during the ONC and the crushed area was 1 mm^2^. However, this crush arrangement being non-standard, the size and topology of the micro-hook unreported and the crush area of the ON assumed rather than measured, their ONC setup and results are difficult to replicate. Some research groups have proposed factory calibrated aneurism clips and other surgical clips delivering fixed crush forces in the range 0.58 N to 1.82 N as a way to standardize ONC models [25–28] because the constant force delivered by these pre-calibrated clips ensures a reproducible injury on each animal, but these clips are not well suited for ONC as their closing forces cannot be controlled and they are difficult to operate, requiring lateral canthotomy surgery to access the ON.

One of the oldest and most common tools employed for ONC are surgical forceps and tweezers. As shown in Table 1, the majority of groups have used a cross-action type design, like the one used in our study here. This design ensures a self-closing action which aids application of a constant force over the crush duration. For example, several groups used Dumont #N5 [15,33,34] or #N7 [16] cross-action tweezers to study the effects of ONC. All of these studies relied on the constant force applied by the self-closing tweezers over various durations to induce graded ON injuries and estimate neurons in the RGC layer of the post-crush ONs. More recently, Huang et al. [35] measured the crushing force between the tips of a cross-action forceps (whose specific make and model was not reported) to be 1.45 N using a lever arm approach, where weights were hung from the tweezer arm tips to hold the tips open, before the ONC. They also estimated the pressure applied by the tips of the forceps on the ON to be 1.93 × 10^6^ Pa based on the forceps tip area of 752 μm^2^.

Although a self-closing, cross-action type tweezers offers the best prospect of delivering a constant crush force on the ON in principle and the crush force can be quantified before the ONC using a method such as the one proposed by Huang et al. [35], the severity of the injury induced by these tweezers in terms of post-crush axon loss still appears to vary from laboratory to laboratory, as is evident from the studies listed in Table 1, all of which have used self-closing tweezers (#N5 or #N7) in mice. Therefore, in this study, we sought to measure the ONC forces in-situ in real time for the first time by sensorizing a Dumont #N7 self-closing tweezers, with dual goals of 1) determining the extent to which crush forces can be controlled by an individual during ONC trials and 2) examining how crush duration and crush force affect RGC survival in ONC trials in mice.

With regards to the first goal, we found that crush forces delivered with a cross-action tweezer by an experimenter during ONC trials, remain nearly consistent (± 8.7% standard deviation) and uncorrelated to the crush duration. This suggests that the variations in the RGC loss reported in ONC studies (Table 1) using self-closing tweezers (#N5 and #N7) could more likely be due to other factors such as the differences between the tip geometries of #N5 and #N7, variations in the crushed areas (due to differences in effective contact widths) of the ONs engaged between the tweezer tips, thus varying the effective crush intensities delivered. With regards to the second goal, we found that axon densities of post-crush ONs closely correlated to the crush impulses delivered, with a survival range of approximately 35% (largest force-impulse) to 75% (lowest force-impulse) axons (Figure 5). Since we only measured the crush forces but not the width of the ON engaged between the tweezer tips (and hence the crush area) in each trial, we did not correlate the injury intensity to injury severity (axon density). Nevertheless, the trend of the relationship between post-crush ON axon density and ONC force-impulse we observed in mice (Fig. 5) is similar to the trend of the relation between the injury intensity to injury severity reported by Huang et al. [35] in rats.

Since the tweezer we employed in this study was a self-closing type, we used the duration of crush time as the primary variable factor to deliver graded force-impulse, and thus, graded injury severity. While this approach to deliver graded injury severity seems most common among researchers studying ONC today, development of a specialized instrument capable of delivering variable crush forces to the ON of any animal model could permit experimenters a much finer control over the extent of axon damage than possible with a self-closing tweezer. Although we cannot conclusively address at this time whether such fine control over the extent of injury would improve nerve crush injury investigations into the mechanisms of neural degeneration and postinjury regeneration, the fact that Duan et al. [4] report differing rates of loss for different retinal ganglion cell types suggests that the ability to control nerve damage with greater precision would likely prove useful. Perhaps an even more important reason to develop an instrument customized for ONC is to simplify and standardize the application of mechanical injury intensity of the traumatic factor (the force applied and the compression sustained by the ON), an important prerequisite related to the quantification of ON injury [35]. Current practice to quantify the injury intensity for forces delivered with a commonly used self-closing tweezer calls for precise measurement of the width of the ON engaged between the tweezer tips and the area of the ON crushed. Such a fine geometric measurement with precision on the order of micrometers during each and every ONC trial is not only challenging but also prone to variations from trial to trial, laboratory to laboratory and model to model. This challenge can be addressed by developing a customized tool for ONC with tips that conform to the geometry of the ON, unlike the sharp, straight or curved tips of typical self-closing tweezers used in ONC studies today.

## Conclusions

In the current study, we investigated the combined effects of crush parameters measured in real time on RGC survival after ON injury in mice using an instrumented, standard self-closing #N7 tweezers. The surviving axon density correlated with crush force, with longer duration and stronger crush forces producing consistently more axon damage. With calibrated instrumented tweezers, like those we employed in this study, experimenters can quantify the actual crush forces applied in situ in real time during ONC to produce consistent and predictable post-crush cell death. The data and knowledge gathered from this study could aid the design and development of a new instrument with geometry and functionality tailored, unlike self-closing tweezers, for ONC experiments to induce consistent damage to the ON with controlled force, duration, and location. Finally, while this study has focused on the crush injury of ONs there is every reason to believe that the findings of this study are applicable to models of nerve crush for other nerves.

## Funding

This work was supported by the National Institutes of Health under Grant number R01EY026286 (X.L.) and Grant number R01EY029121 (X.L.); and the National Science Foundation under Grant number 0938072 (L.S.).

## Notes

#### Summary of Updates

edited abstract.

